# Whole genomes reveal severe bottleneck among Asian hunter-gatherers following the invention of agriculture

**DOI:** 10.1101/2020.06.25.170308

**Authors:** Saikat Chakraborty, Analabha Basu, the GenomeAsia 100K Consortium

## Abstract

The invention of agriculture (IOA) by anatomically modern humans (AMH) around 10,000 years before present (ybp) is known to have led to an increase in AMH’s carrying capacity and hence its population size. Reconstruction of historical demography using high coverage (~30X) whole genome sequences (WGS) from >700 individuals from different South Asian (SAS) and Southeast Asian (SEA) populations reveals that although several present day populous groups did indeed experience a positive Neolithic Demographic Transition (NDT), most hunter-gatherers (HGs) experienced a demographic decrease. Differential fertility between HGs and non-HGs, exposure of HGs to novel pathogens from non-HGs could have resulted in such contrasting patterns. However, we think the most parsimonious explanation of the drastic decrease in population size of HGs is their displacement/enslavement by non-HGs.

**Significance Statement:** The invention of agriculture, around 10000 years ago, facilitated more food production which could feed larger populations. This had far-reaching socio-political and demographic impacts, including a ~10,000 fold increase in global population-size in the last 10,000 years. However, this increase in population size is not a universal truth and present day hunter-gatherer populations, in contrast, have dwindled in size, often drastically. The signatures of this rise in population size are discernible from the genomes of present-day individuals. Using genomic data, we show that for the majority of Asian hunter-gatherers, population-sizes drastically decreased following the invention of agriculture. We argue that a combination of displacement, enslavement and disease resulted in the decimation of hunter-gatherer societies.

## Introduction

Archaeological and anthropological evidence suggest that the invention of agriculture was a protracted process that developed through “gathering”, “cultivation” and “domestication” and took several millennia to be standardized. During these initial stages populations were not exclusively gatherers or cultivators but moved freely among different lifestyles across seasons or years and utilized a mix of these different lifestyle strategies (*1–3*). Between 11,000 to 8000 years before present (ybp), almost all the major present day cereal crops were domesticated in different parts of Asia (*4*). The invention of agriculture (IOA) is believed to have been a major event in the history of anatomically modern humans (AMH) as it generated enough surplus to have given rise to the state and its associated machinery, particularly division of labor – producers and non-producing elites, and possibly armies (*2*). An immediate consequence of IOA was an increase in the carrying capacity of AMH and therefore its population size. This has been termed as the Neolithic Demographic Transition (NDT) (*5*). However, there is accumulating Archeological and Anthropological evidence that the impact of IOA was not uniform for all societies. Not all populations experienced an increase in carrying capacity. Neither did they readily transited from a hunter-gatherer (HG) way of life to an agrarian society (AS) even after being exposed to the methods of cultivation (*2, 3*). In fact an enormous number of indigenous populations still are HGs. These groups are not expected to experience an increase in their population size, unlike populations that took up agriculture as their primary mode of livelihood. In this paper we contest the popular connotation of NDT and ask the question - How the demography of present day hunter-gatherers were affected following the invention of agriculture?

We tried to answer the question using the whole genome sequences (WGS) for South and Southeast Asian (SAS and SEA) populations, both HGs and non-HGs, generated as part of the GenomeAsia100k Consortium. SAS and SEA together are home to ~1/3 of the current global AMH population (*6*). The methods of modern agriculture were introduced quite early in these parts of the world (*7*). However, several indigenous SAS and SEA populations still practice a HG way of life or used to be HGs even a few generations ago. Such populations are not expected to go through a demographic transition, although a secondary transition to being HG from AS during the early days of IOA cannot always be ruled out.

Genomes sequences contain a vast amount of information about our historical past. This information can be gleaned using appropriate theoretical and computational approach. Coalescent theory is the cornerstone of modern population genetics that tries to understand population genetic processes by constructing gene genealogies and estimating time to most recent common ancestor (TMRCA) for *a* genetic variant site in a sample. The rate at which genealogies reach TMRCAs, also known as the coalescent rate, varies with population size. Contracting and expanding population lead to higher and lower coalescent rates (CR), relative to CR in a population with constant size. Therefore, given the right kind of genetic information, this framework can be used to infer demographic histories of populations. Coalescent modelling using WGSs from present day individuals is by far the most robust method to reconstruct demographic histories. Two such widely used methods are Pairwise and Multiple Sequential Markovian Coalescnet (PSMC and MSMC respectively) (*8, 9*). In case of PSMC, only one diploid WGS can be used at a time. Although, several samples can be used in case of MSMC, it is extremely computationally intensive. In this work we have employed a method, *smc*++ that uses genetic information from all the individuals from a specific population (*10*). Using this method, we have inferred the demographic histories of 59 sociologically defined populations (Details in **Table S1**) across SAS, SEA and Oceania (OCE) along with one each from Africa (YRI), Europe (GBR) and one Northeast Asia (HAN) to test if the HGs and ASs show similar demographic pattern around the time of IOA. We have found that while the effective population sizes (N_e_) of almost all non-HG populations increased around 10,000 years before present (k ybp), most Asian HG populations experienced severe population bottleneck during the same period.

## Results

We carried out extensive coalescent modelling for 59 sociologically defined populations (Details in **Table S1**) across SAS, SEA and Oceania (OCE) along with one each from Africa (YRI), Europe (GBR) and one Northeast Asia (HAN) to test if the HGs and ASs show similar demographic pattern around the time of IOA. We used the Sequential Markovian Coalescent Prime (SMC’) algorithm (*11*) as implemented in the program *smc*++ (*10*) to infer the historical demographics since 500k years before present (ybp) from whole genome sequences (WGS) (**Fig. 1**). We took generation time as 25 years.

**Fig. 1.**
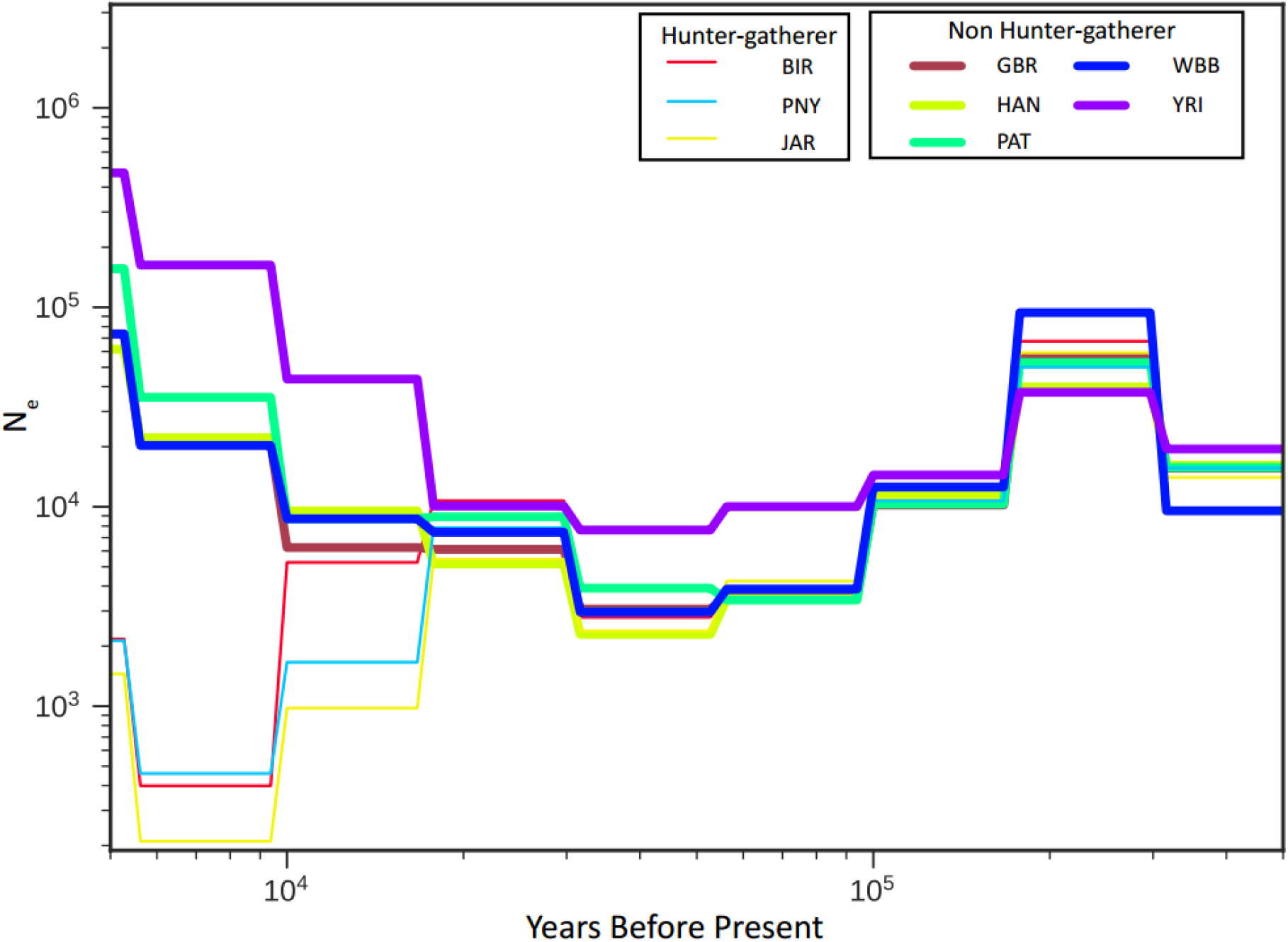
Demographic histories of selected populations used in this study. Effective population size (N_e_) change is shown for the period 500k to 5k years before present (ybp) The X and Y axes are in *log*_*10*_ scale. YRI, GBR and HAN refer to Yoruba, Great Britain and Han Chinese respectively (See **Supplementary Table S1**) and are included here as representatives of Africans, Europeans and Asian AMH populations. As has been previously shown, the non-African populations separate out from the African YRI ~100k ybp. These non-African populations experienced a strong bottleneck ~60k ybp which they start recovering from ~30k ybp. Around 10k ybp, along with GBR and HAN, the Pakistani population Pathan (PAT) and the upper caste Hindu group West Bengal Brahmin (WBB) show an increase in N_e_, while the hunter-gatherer (HG) SAS groups Paniya (PNY), Birhor (BIR) and Jarwa (JAR) show a sharp decline. This departure approximately coincides with the time when agriculture was independently invented in different parts of Asia.

The general pattern observed by us until NDT is similar to other works that have used WGS and SMC’ or related methods to infer demographic history (*8, 9*). The non-African populations show separation from the Yorubans (YRI) ~100k ybp followed by a strong bottleneck ~60k ybp, a testimony to the ‘out-of-Africa’ migration (*8,9*). However, we observe an increase in the effective population size (N_e_) for the Great Britain (GBR), Han Chinese (HAN), the upper caste West Bengal Brahmins (WBB) and the Pathans (PAT) ~10k ybp coinciding with IOA. During the same time, a sharp decline in N_e_ for the SAS HG groups Birhor (BIR), Paniya (PNY) and Jarwa (JAR) was observed [Details of the populations in **Table S1**].

We extended the analysis to other populations inhabiting SAS, SEA and Oceania (OCE) and investigated their demographic histories. This dataset consisted of 724 individuals from 59 populations, which we grouped based on geography, language and lifestyle (Hunter-gatherers (HGs) and Non-hunter-gatherers (non-HGs)). We define as HGs populations i) that live by hunting and gathering, ii) that use a mix of subsistence, including slash and burn, cultivation along with hunting and gathering and iii) pastoralists. We first investigated if there were overall difference in N_e_ between HGs and non-HGs around IOA in this entire region. We calculated N_e_ at ~5.6k ybp, 10k ybp and ~17.8k ybp for each of the population and compared the median N_e_s of HGs and non-HGs across 1000 bootstrap replicates (**Fig. 2B**, **Table S2**). We observed no difference between the two groups before IOA (at ~17k ybp). However, this difference was significantly higher ~10k ybp (non-HGs>HGs) and this difference was maintained at least for the next five millennia. We also carried out this comparison only for the SAS populations and observed similar patterns. However, the difference was not significant between ~17k ybp and the two recent timepoints (**Fig. S1** and **Table S3**). We further estimated the growth rate for these populations from 10k ybp to ~5k ybp assuming an exponential growth. Again we observed considerable heterogeneity among the populations in each lifestyle classes (HG or Non-HG) (**Fig. 2C**). Moreover, although the point estimates for bootstrapped medians were different, there were slight overlap between the upper CI for HGs and the lower CI for the Non-HGs (Median(Upper CI, Lower CI): HG = 0.992 (0.994, 0.990); Non-HG = 0.998 (1.004, 0.993), **Fig. 2D**). This suggests that significant difference between the N_e_s can arise even if there is a small difference in the overall growth rates of the populations, conditional on the assumption of long-term continuous exponential growth/decline, in the two classes. We could model the growth rates from 10k to 5.6k ybp. It is clear from the demography figures (**Fig. 1** and **Fig. S2-6**) that for most populations the transition (increasing or decreasing N_e_) began a little before 10k ybp. Therefore, these reported growth rates are either underestimates (for increasing transition) or overestimates (for decreasing transition).

**Fig. 2.**
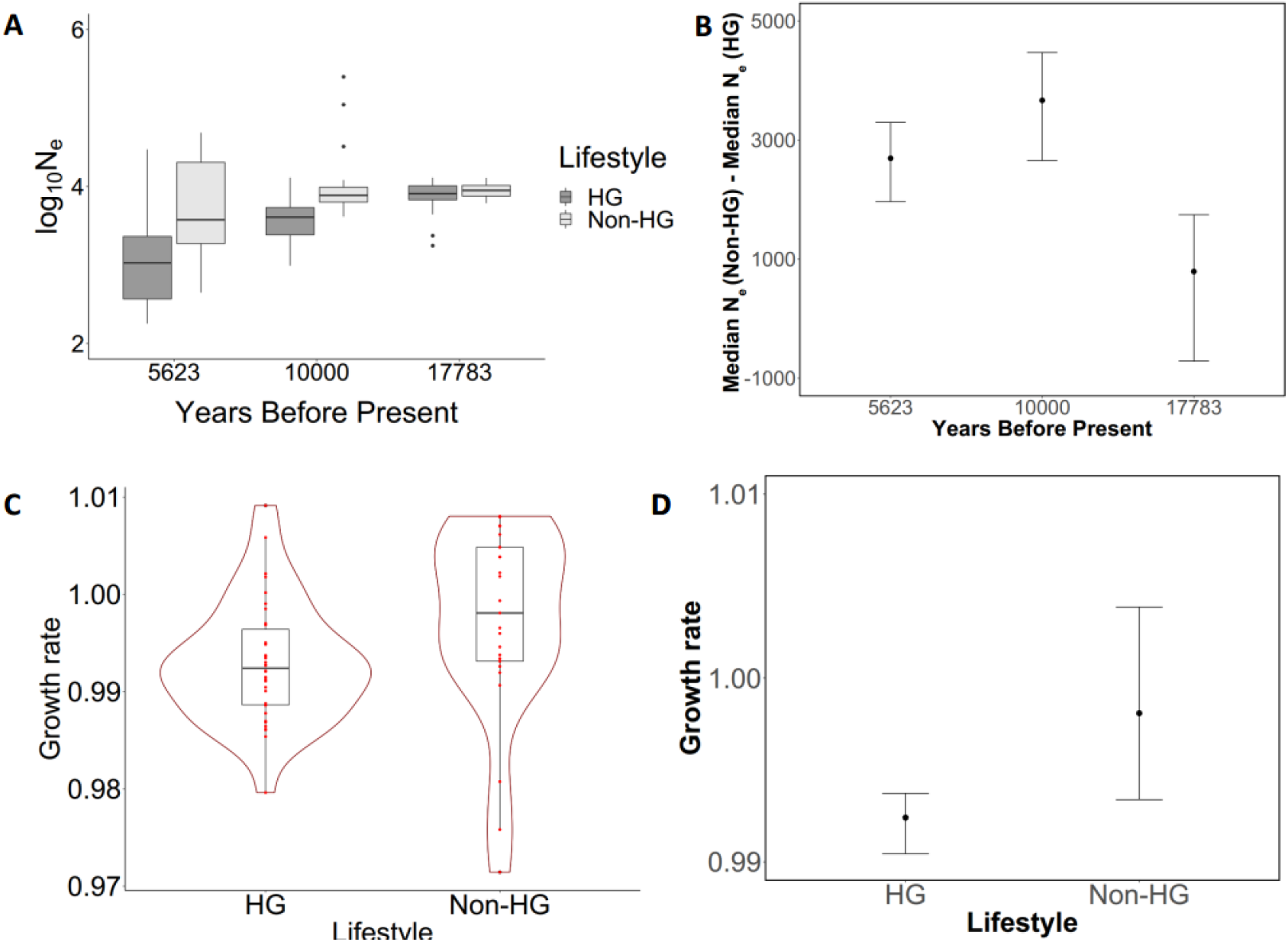
Comparison of HG and Non-HG demography. **A.** Distribution of effective population size (N_e_) between hunter gatherers (HG) and non-hunter gatherers (Non-HG) at three time points obtained from the coalescent model spanning before and after 10,000 years before present (10k ybp). **B.** Median and 95% confidence interval (CI) of the median N_e_ difference between HGs and non-HGs across 1000 bootstrap replicates. The CI at 17783 ybp does not overlap with the CIs at 5623 and 10000 ybp. However, there is overlap between the CIs at the two more recent timepoints (N_HG_=34; N_Non-HG_=25). **C.** Distribution and median of growth rate assuming exponential growth since 10k ybp for different lifestyles. The median and 95% CI across 1000 bootstrap medians are plotted in figure **D.** Slight overlap between the upper CI of HGs and lower CI of Non-HGs is observed. The populations used in this comparison are marked with a * in **Table S1.**

We further investigated this pattern by focusing on each of the three regions, i.e., SAS, SEA and OCE. We grouped the SAS populations based on geography or language, as both of these have been shown to have substantial explanatory power for genomic variation in SAS populations (*12*).In our dataset, the most widely spoken languages by SAS HGs belong to the Dravidian (DR) language family. A large number of ‘upper’, ‘middle’ and ‘lower’ caste Hindus living in southern India also speak in DR family languages. However, Austroasiatic (AA) language speakers, inhabiting middle and eastern India today are almost exclusively HG tribes. The people in northern and eastern India, Pakistan and Nepal, almost exclusively speak in Indo European (IE) language, and the people inhabiting northeastern India speak languages belonging to the Sino-Tibetan (ST) language family. The populations living in the Andaman and Nicobar Islands speak in the isolated Ongan which is speculated to belong to the AA language family.

We observed that the aboriginal HG populations Jarwa (JAR), Onge (ONG) and Nicobarese (NIC) inhabiting the Andaman and Nicobar Islands show two distinct population bottlenecks, ~20k ybp and ~10k ybp (**Fig. S2A**). This bottleneck ~20k ybp roughly coincides with the deglaciation following the last glacial maximum and may point towards local migration and/or isolation (*13, 14*). In case of the Austroasiatic speaking HG tribes from mainland SAS, in concordance with the islanders, we observed a pronounced bottleneck ~10k ybp, but unlike the islanders, we did not observe a population bottleneck ~20k ybp (**Fig. S2B**). We observed considerable heterogeneity among the Indian DR tribes (**Fig. S3**). Out of the 12 DR speaking tribes in our dataset, seven went through a bottleneck ~10k ybp (**Fig. S3B**), while five did not (**Fig. S3C**). These seven bottleneck experiencing tribes did not have characteristically lower F_st_ with the AA tribes than those that did not go through bottleneck (**Fig. S7**). [The Pakistani DR speaking tribe Brahui (BRA) is discussed later with other Pakistani tribes.] There are very few HG tribes among the IE speakers. However, ‘lower’ caste IE speakers have substantial SAS HG ancestry (*15*).Three out of four IE speaking ‘lower’ castes (Bagdi (BAG), Chamar (CHM) and Nababuddha (NAB)) we studied showed a decline in N_e_ from ~20k ybp onwards, while the fourth, Mahar (MHR), increased during the same time (**Fig. S4A**). Among the Indian ‘upper’ castes, Iyer (IYE), West Bengal Brahmin (WBB) and Saurashtra Brahmin (SRB) showed an increasing N_e_ while the rest (Iyengar (IYA), Gaud (GAU), Soryupari Brahmin (SOB) and Agharia (AGH)) experienced a decreasing trend ~10k ybp (**Fig. S4B**). Among the 7 populations potted in **Fig. S4B**, IYE, IYA and SOB are DR speakers, while the rest speak in IE languages. Except Brahui (BRA) and Hazara (HAZ), N_e_ for all other Pakistani populations either increased or remained constant during this period. The Brahui and Hazara populations experienced a marked increase in their N_e_ ~20k – 10k ybp followed by a bottleneck ~10k ybp (**Fig. S5A**). It must be noted here that Brahui is an isolated DR language speaking population from Pakistan. All the other populations from this region in our dataset speak in IE language. However, historically Hazaras are known to have spoken in a Mongolic family language (*16*) and their current population is admixed between Ancestral North Indian (Pathan like) and Ancestral Tibeto-Burman (Mog like) ancestries (*12, 15*). Among the five Sino-Tibetan speaking SAS populations for which we inferred demographic histories, Mog (MOG) did not go through a bottleneck ~10k ybp, but the others did (Chakma (CHK), Toto (TTO), Jamatia (JAM) and the ‘upper’ caste Manipuri Brahmin (MNP)). The bottleneck was most severe for the endangered tribe Toto (**Fig. S5B**).

In case of the Southeast Asian populations (**Fig. S6**), we observed only two populations, namely Kinh (KIN) and Dai (DAI), among the 11 for which we estimated demographies, to have experienced an increase ~10k ybp (**Fig. S6A**). The Kinh people are the most numerous among the present day ethnic groups inhabiting Vietnam. In fact their language, i.e., Vietnamese, is the most widely spoken Austroasiatic language in the world (*17*). On the other hand, there are >1 million Dai people currently distributed in southwestern China and Myanmar (*18*). The N_e_ for Rampasasa ‘pygmies’ (RAM) of Indonesia and the Phillipine Ati (ATI) people, did not change around 10k ybp (**Fig. S6C**). Both RAM and ATI speak in Austronesian languages and lead a HG lifestyle. All the rest of the 7 populations experienced a bottleneck ~10k ybp (**Fig. S6D**). These are the Malaysian populations Senoi Semai (SNS), Temuan (TEM), Kensiu (KEN), Kintak (KIN), and from Indonesia the Austronesian group NIA, the Cibal (CIB) and the Bena (BEN) people. The NIA, CIB, BEN, RAM, TEM and ATI people speak in Austronesian, while the rest, i.e., SNS, KIN and KEN people speak in Austroasiatic languages.

All of the four Oceanian populations in our dataset, namely, Nakanai (NAK), NKB (Nakanai Bileki), Baining (BAI) and Papuan (PAP) were observed to have experienced bottleneck ~10 k ybp (**Fig. S6B**). Among these the BAI people are small subsistence farmers and the rest are HGs. The language of the BAI people occupy a family of their own, whereas the NAK and NKB speak in Austroasiatic languages, and the language of the PAP belong to the Papuan language family.

## Discussion

We observed 27 out of the 34 hunter-gatherers across SAS, SEA and OCE to have experienced severe bottleneck following IOA. Most HGs that experienced a decrease in N_e_ during the same time inhabit isolated locations as small populations. The populations from the Andaman and Nicobar Islands, i.e., Jarwas, Onges and Nicobarese, and that from the Flores island such as the Bena people and the Rampasasa pygmies are restricted to their island homes and are naturally isolated and small. But, even the Dravidian and Austroasiatic speaking tribes from mainland India, e.g., Birhors, Paniyas, Oraons, Mundas, etc., and from mainland Malaysia e.g., Senai, Kintak, Kensiu, etc., all have present distribution restricted to isolated forested highlands or plateaus that are not suitable and/or fertile enough for extensive agriculture. Many of these populations, however, practice some form of subsistence including slash-and-burn cultivation.

The primary reason suggested for NDT to have taken place is sedentism (*2, 19*). Agriculture forced people to become sedentary resulting in an increase in the rate of reproduction. The crowding associated with a sedentary lifestyle also led to an increase in mortality. However, the overall fertility, i.e. *difference between the number of births and deaths,* were more among ASs than among HGs (*2*). This can happen because of the following. 1) A carbohydrate rich diet, instead of a diet dominated by seeds, nuts and animal protein, enabled early weaning of the children. Consequently, AS women became fertile sooner following childbirth than HG women, resulting in smaller inter-offspring interval among the former. 2) The cereal dominated diet of the AS people led to an earlier onset of menarche and later onset of menopause than the high protein diet of the HG people, resulting in a longer reproductive period among the ASs. 3) As travelling with and taking care of more than two children at a time is virtually impossible in case of a nomadic lifestyle, the HGs might have even practiced infanticide, abortion and sexual abstinence to keep the number of children in the group within limits (*3, 19, 20*).

Although, these explain the increase in N_e_ while transiting from a primarily hunter-gatherer to a primarily settled agriculturist lifestyle, it does not explain why N_e_ is case of most HGs should decrease following IOA. At the most one would expect the HG population size to remain unchanged. Here we discuss our results in light of two mutually non-exclusive lines of evidence in the literature. Domestication of plants and animals led to major changes in the kind of environment hitherto experienced by AMHs. Increase in the size and density of AMH population in the early settlements also resulted in an increase in the size and density of the domesticated animals and plants and their associated pests and parasites. This facilitated opportunities for host jumping exposing AMHs to a plethora of novel infections they had not experienced till that point, resulting in pandemics and epidemics. The high crowding of people, animals and pests in Neolithic settlements also ensured fast transmission of these pathogens. Many early and late Neolithic sites were rapidly abandoned for no apparent reason. Although it is not clear why this might have happened, the most plausible reason could have been epidemics (*2*). It has been argued that acute community infectious diseases affecting AMH such as influenza, measles, mumps, smallpox, etc. that are characterized by low incubation time, short life cycle, lack of intermediate host, high contagiousness and seasonality, are result of zoonosis from livestock (*3*). To be maintained they require population sizes of the order that could only exist in settled agriculturist societies. (*21*) Such diseases, it has been argued, did not exist in pre-agricultural societies and are still rare in isolated hunter-gather societies (*22*). Albeit the significant fitness cost, large and dense populations allow both the host population to evolve resistance or tolerance towards the disease as well as leads to less virulence of the pathogens (*23*). Therefore, the consequences of such epidemics are more severe in small isolated populations than in large populations as has been recorded in case of the Andaman islands (*24, 25*), Caribbean islands (*26*), Hawaii (*27*) and other parts of the new world and Oceania (*3, 28, 29*).

The second reason for observing contrasting demographic transition between HGs and non-HGs could be mass enslavement and displacement, if not killing, of HGs by ASs (*2, 3*). Agriculture is extremely labor intensive and producing enough to eat for everyone in the dense and large early settlements would have required a huge workforce. Moreover, early domesticated crops, although higher yielding compared to their wild relatives, were neither as productive as modern cultivars nor were they adapted to largescale farming (*30*). Therefore, producing enough to feed an ever growing population would require enormous amount of labor (*2*). Consequently, the division of labor and formation of the non-producing elites by the time the early city states came into being also simultaneously gave rise to a huge population of bonded-laborers. Historical evidence suggests that people, often HGs, that were defeated in war constituted the major proportion of this labor force (*31*). In fact, such slaves formed the main booty of the war. Moreover, selectively women and children were enslaved more frequently and the men either got killed or fled away (*2*). One may extend such phenomena only a few millennia back in time and argue that enslavement was being practiced even during pre-historical times (*32*). The advantage in the number of the AS people facilitated subjugation of the HGs and thereby increases their workforce through slave laborers. A huge number of HG people may also have been either killed or displaced during this process as is known to have happened more recently in Southeast Asia (*2, 33*). This resulted in the depletion of the HG populations and hence low N_e_. The decrease in females and young children meant skewed sex ratio and inverted age pyramid further lowering the N_e_. The enslaved HG people, on the other hand, would have become naturalized into the ASs within a few generations and leading to increased AS N_e._

Therefore a combination of a) subjugation in the form of abduction and enslavement followed by assimilation of HGs by ASs, b) widespread killing and displacement of HGs by ASs and c) decimation of HGs in the face of parasites introduced by ASs, might have resulted in the difference in N_e_ between HGs and ASs following IOA (*2, 3, 23, 33*). The fact that most populations still practicing a HG lifestyle are cornered and inhabit small isolated populations in relatively uninhabitable habitats such as forests, mountains, arid plateaus and deserts further supports this plausible explanation. Although historical and anthropological evidence supporting the importance of the above mentioned processes in bringing about differences in demography and lifestyle during the formation of early states are mounting, here we provide genetic evidence to show that such changes began as soon as AMHs began practicing settled agriculture, long before the first city states came into being.

## Materials and Methods

### Data

All the data included in this study are from the Genome Asia 100K Consortium dataset (GA100K). Out of the 211 populations included in GA100K, we selected 62 to infer the finer demographic patterns of South Asia (SAS), Southeast Asia (SEA) and Oceania (OCE) and for comparison with populations from Northeast Asia, Africa and Europe. The details of the populations used in this study are tabulated in **Table S1**.

### Coalescent modelling

In order to infer the population size change histories or demographic histories (DH), we used the program *smc*++ (*10*). The master GA100k *vcf* file was used to generate population specific input files. The GRch37/hg19 ‘Gap’ tracks downloaded from the UCSC table browser were used as a mask file (The `−m` flag) (*34*). An example of the command used to generate the input file is shown (for the population MOG) below.

~~~
for i in $(seq 1 22); do
smc++ vcf2smc substitutions.annot.vcf.gz
/home/users/ntu/nidhan/scratch/smcpp_masked/MOG/mog.chr“$i”.masked.sm
c.gz “$i” -m
/home/users/ntu/nidhan/scratch/smcpp/mask_files/hg19_gaps.sorted.bed.
gz
MOG:GA001020,GA001021,GA001022,GA001023,GA001024,GA001025,GA001026,GA
001027,GA001028,GA001029,GA001030,GA001031,GA001032,GA001033,GA001034
,GA001035,GA001037,GA001038,GA001039,GA001040 --cores 24
done
~~~

DH was estimated using the Powell’s algorithm (*9*) between 40 to 40000 generations. Mutation rate was taken as 1.25×10^−8^ (*10*). All estimations were performed taking 12 knots. An example command (for the population HAZ) is shown below.

~~~
smc++ estimate --cores 24 --algorith Powell --knots 12 --timepoints
40,40000 -o /home/users/ntu/nidhan/scratch/smcpp_masked/HAZ_12knots
1.25e-8 /home/users/ntu/nidhan/scratch/smcpp_masked/HAZ/*.smc.gz
~~~

The model JavaScript Object Notation (*json*) files were plotted from 5k to 500k ybp using the `plot` command in *smc*++. Generation time (`−g` flag) was taken as 25 years. A sample is shown below.

~~~
smc++ plot BUR.pdf -g 25 -x 5000 500000 -y 100 1000000
model.final.json
~~~

### F_st_ estimation

We estimated pair-wise F_st_ between Austroasiatic (AA) and Dravidian (DR) speaking tribes. The *vcf* file was converted into a PLINK(*35*) format file and data for the 17 AA and DR tribal individuals were filtered out. This was further converted into EIGENSTRAT format using unix shell commands and the CONVERTF program. The pair-wise F_st_ values were estimated using the *smartpca* program included in the EIGENSTRAT package (*36*). The results are shown in Fig. S9 all pairs except Brahui (BRA) for reasons mentioned in the main text and Toda (TOD) as the DNA samples sequenced were not of good quality.

### Statistical analyses

We compared the N_e_ change before, during and after the Neolithic revolution between hunter gatherers (HG) and non-hunter gatherers (Non-HG) using bootstrap resampling. For each time point difference of the median of N_e_ between HGs and non-HGs was calculated. This was repeated 1000 times to get a distribution of median differences. The median of this distribution at each time point along with its 95% confidence interval is shown in **Figure 2B**. The populations that were included in this analysis are marked with an ‘*’ in **Table S1** and the results of the bootstrap analysis is summarized in **Table S2**. The effective population sizes (N_e_) at three time points spanning 10k ybp was calculated from the *smc*++ model outputs. *smc*++ uses a 2N_0_ scaling. Therefore the X axis was taken as *2*\**N*_*0*_\**(model knot value)* and the Y axis was taken as *N*_*0*_*\*exp(model y value)* [because *(model y value)*=*log(N*_*t*_/*N*_*0*_)]. The three time points from the model were 5623k ybp, 10000k ybp and 17783k ybp. The function *sample* in *R* (*37*) was used for generating the bootstrap samples (*replace* = *T*). Growth rate (R) for each population was estimated by fitting the equation *R*=*(N*_*t*_/*N*_*0*_)^*1/t*^, where *N*_*t*_ is the *N*_*e*_ at 5623 ybp (225 generations before present (gbp)), *N*_*0*_ is *Ne* at 10000 ybp (400 gbp) and *t* for all cases is (400-225) = 175 generations. We generated bootstrap samples for medians separately for HGs and Non-HGs as described above and plotted the median and 5% CIs for each distribution. Plotting was performed using *ggplot2 (38).*

## Supplementary Materials

Fig. S1. Comparison of SAS HG and Non-HG populations across the three time points around IOA.

Fig. S2. Demography of populations inhabiting SAS islands and those speaking in Austroasiatic family languages.

Fig. S3. Demography of Dravidian language speaking indigenous tribes.

Fig. S4. Demographic histories of Indian Indo-European speaking caste populations.

Fig. S5. Demographic histories of Pakistani populations and those speaking in Sino-Tibetan family languages.

Fig. S6. Demographic histories for SEA and OCE populations included in this study.

Fig. S7. F_ST_ estimates for the Indian DR and AA speaking tribes.

Table S1. Description of the populations used in this study.

Table S2. Summary of the bootstrap results.

Table S3. Summary of the bootstrap results for SAS populations.

## Additional Data

Data S1. GenomeAsia 100K Consortium Members

## General

The authors would like to thank Anasuya Chakrabarty and Diptarup Nandi for help with the analysis of data, and Anasuya Chakrabarty, Diptarup Nandi, Debashree Tagore, Jeffrey D Wall and Partha P Majumder for helpful comments and suggestions. The data used in this work were generated as part of the GenomeAsia 100K Consortium. Most of the computational work was performed using the facilities of the National Supercomputing Centre, Singapore.

## Author contributions

SC and AB conceptualized the project. SC performed the computational and statistical analysis. AB interpreted the results. SC and AB co-wrote the manuscript. AB supervised the project.

## Data and materials availability

GeomeAsia 100k individual VCF files are available from the European Genome-phenome Archive (EGA) under accession ID EGAS00001002921. The csv files and R codes can be obtained from the following repository on request https://github.com/saikat-popgen/Coalescent_paper

