## Supplementary File for "Whole genomes reveal severe bottleneck among Asian hunter-gatherers following the invention of agriculture"

**GenomeAsia 100K Consortium**
Jeffrey D. Wall, Eric W. Stawiski, Aakrosh Ratan, Hie Lim Kim, Changhoon Kim, Ravi Gupta, Kushal Suryamohan, Elena S. Gusareva, Rikky Wenang Purbojati, Tushar Bhangale, Vadim Stepanov, Vladimir Kharkov, Markus S. Schröder, Vedam Ramprasad, Jennifer Tom, Steffen Durinck, Qixin Bei, Jiani Li, Joseph Guillory, Sameer Phalke, Analabha Basu, Jeremy Stinson, Sandhya Nair, Sivasankar Malaichamy, Nidhan K. Biswas, John C. Chambers, Keith C. Cheng, Joyner T. George, Seik Soon Khor, Jong-Il Kim, Belong Cho, Ramesh Menon, Thiramsetti Sattibabu, Akshi Bassi, Manjari Deshmukh, Anjali Verma, Vivek Gopalan, Jong-Yeon Shin, Mahesh Pratapneni, Sam Santhosh, Katsushi Tokunaga, Badrul M. Md-Zain, Kok Gan Chan, Madasamy Parani, Purushothaman Natarajan, Michael Hauser, R. Rand Allingham, Cecilia Santiago-Turla, Arkasubhra Ghosh, Santosh Gopi Krishna Gadde, Christian Fuchsberger, Lukas Forer, Sebastian Schoenherr, Herawati Sudoyo, J. Stephen Lansing, Jonathan Friedlaender, George Koki, Murray P. Cox, Michael Hammer, Tatiana Karafet, Khai C. Ang, Syed Q. Mehdi, Venkatesan Radha, Viswanathan Mohan, Partha P. Majumder, Somasekar Seshagiri, Jeong-Sun Seo, Stephan C. Schuster & Andrew S. Peterson

**This file includes:**

Figs. S1 to S7
Tables S1 to S3

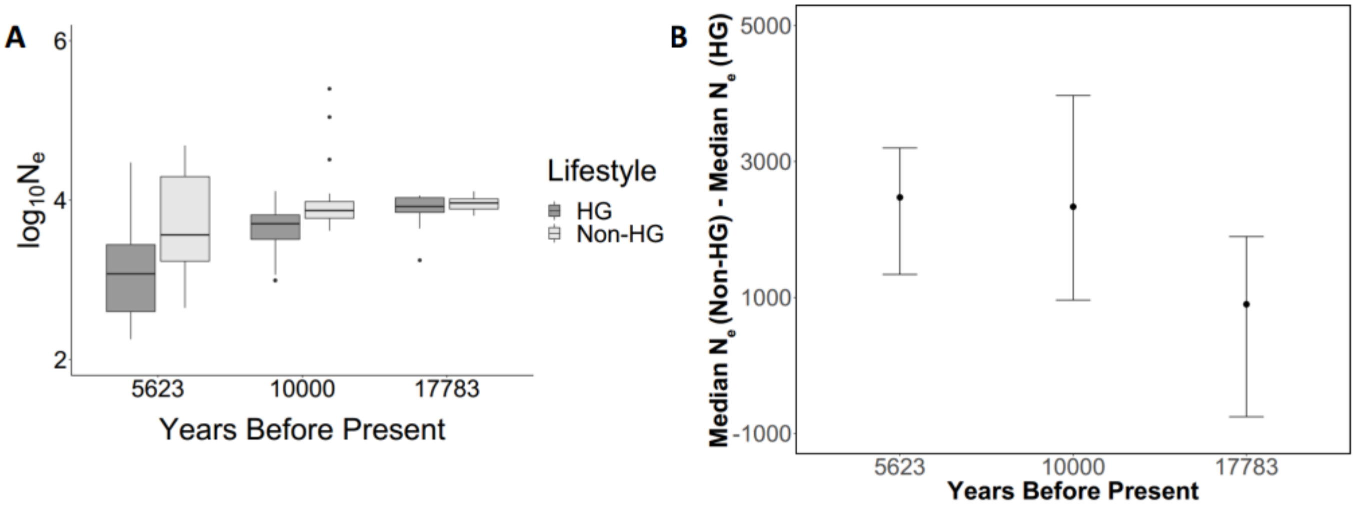


**Fig. S1.**

**Comparison of SAS HG and Non-HG populations across the three time points around IOA. A.** Distribution of effective population size ($N_e$) between SAS hunter gatherer (HG) and non-hunter gatherers (Non-HG) at three time points obtained from the coalescent model spanning before and after 10,000 years before present (10k ybp). **B.** Median and 95% confidence interval (CI) of the median $N_e$ difference between HGs and non-HGs across 1000 bootstrap replicates. Unlike for the entire SAS, SEA and OCE dataset , the upper CI at 17783 ybp does overlap with the lower CIs at 5623 and 10000 ybp ($N_{HG}$=21; $N_{Non\text{-}HG}$=23).

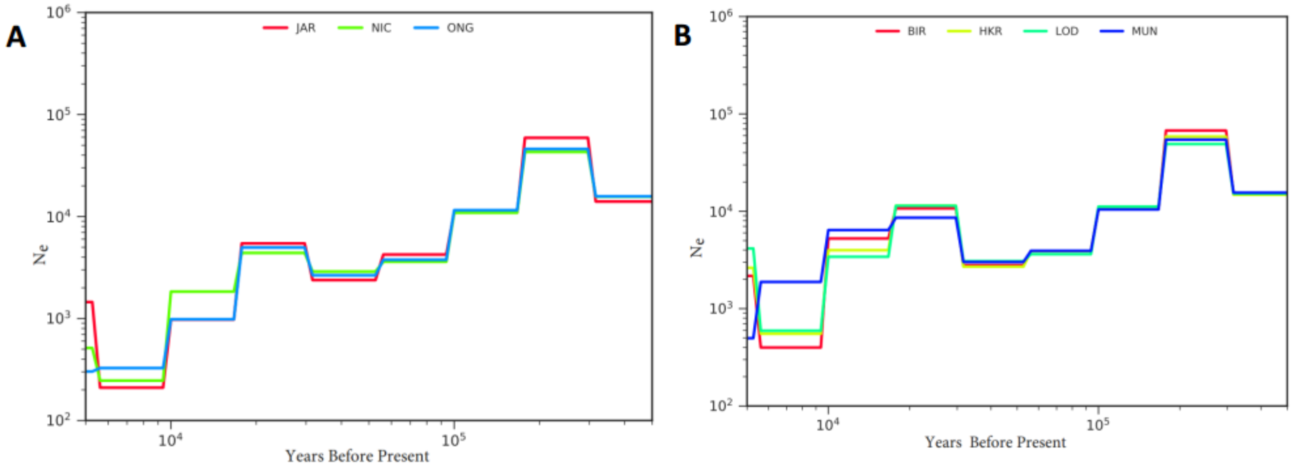


**Fig. S2.**

**Demography of populations inhabiting SAS islands and those speaking in Austroasiatic family languages. A.** Demographic histories for the populations inhabiting the Andaman and Nicobar Islands. Population size change histories are plotted from 500k ybp to 5k ybp. The populations are Jarwa (JAR), Onge (ONG) and Nicobarese (NIC). Please refer to **Table S1** for the details of these populations. **B.** Demographic histories for mainland South Asian Austroasiatic speaking tribes. Population size change histories are plotted from 500k ybp to 5k ybp. The populations are Birhor (BIR), Hill Korwa (HKR), Lodha (LOD) and Munda (MUN). Please refer to **Table S1** for the details of these populations.

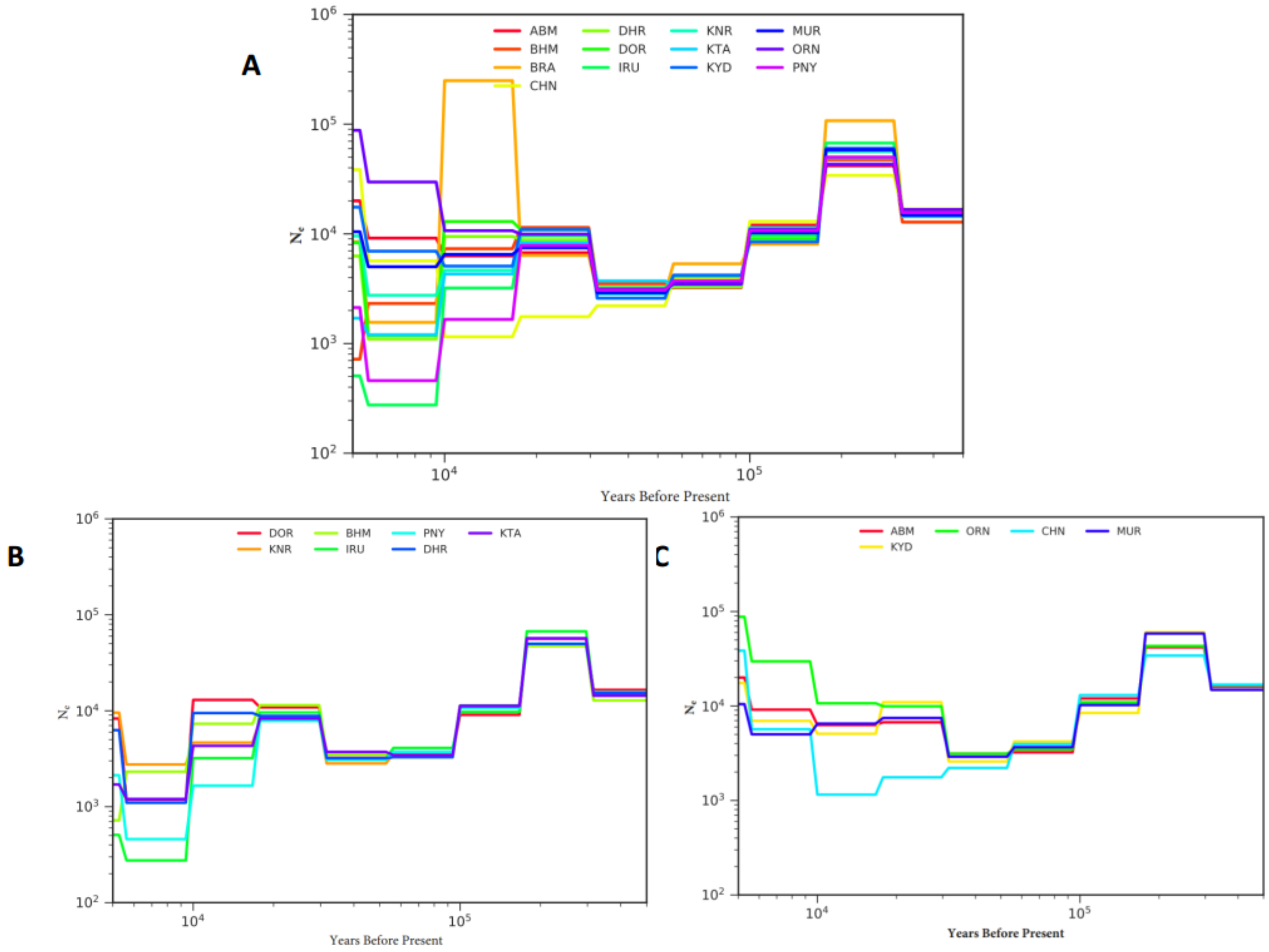


**Fig. S3.**

**Demography of Dravidian language speaking indigenous tribes. A.** Demographic histories for mainland South Asian Dravidian speaking tribes. Population size change histories are plotted from 500k ybp to 5k ybp. The populations are Abujmaria (ABM), Chenchu (CHN), Irula (IRU), Muriya (MUR), Bison Horn Maria (BHM), Dhurwa (DHR), Kondareddi (KNR), Kota (KTA), Oraon (ORN), Brahui (BRA), Dorla (DOR), Kaya Dora (KYD) and Paniya (PNY). The populations that went through a bottleneck ~10k ybp are shown in Figure **B** and the populations that experienced no change or an increase in Ne are shown in Figure **C**. BRA has been excluded from Figure B because of its peculiarities. (See main text for details). Please refer to **Table S1** for the details of these populations

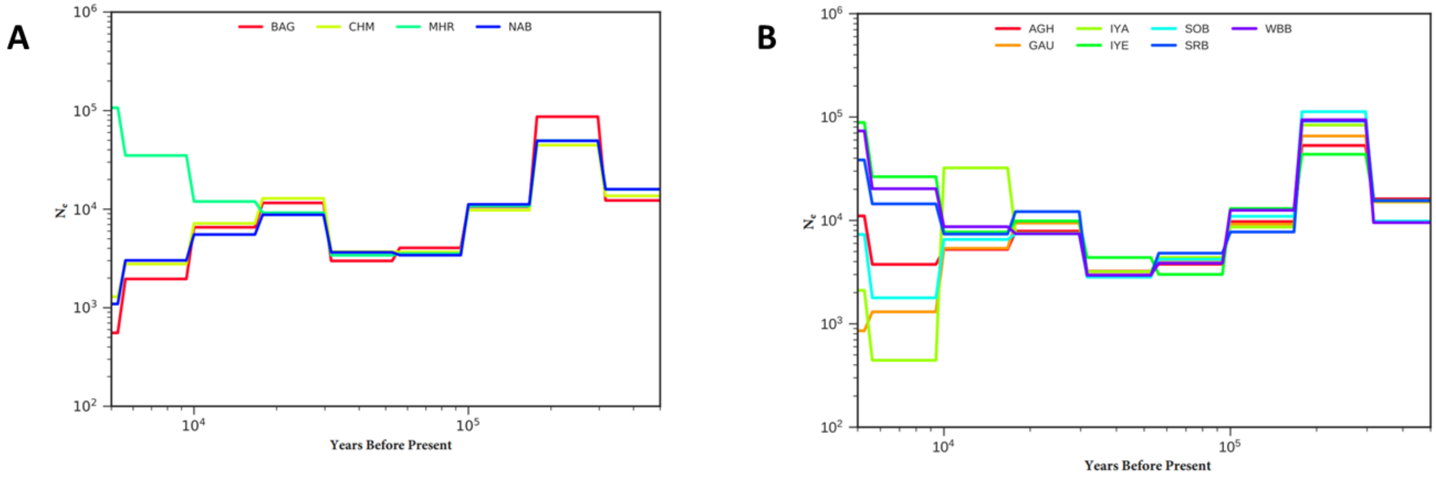


**Fig. S4.**

**Demographic histories of Indian Indo-European speaking caste populations.** The DH for ‘lower’ and ‘upper’ caste populations are shown in Figures **A** and **B** respectively. Population size change histories are plotted from 500k ybp to 5k ybp. The populations are Bagdi (BAG), Chamar (CHM), Mahar (MHR), Nababuddha (NAB), Agharia (AGH), Iyengar (IYA), Saryupari Brahmin (SOB), West Bengal Brahmin (WBB), Gaud (GAU), Iyer (IYE), Saurashtra Brahmin (SRB). Please refer to **Table S1** for the details of these populations.

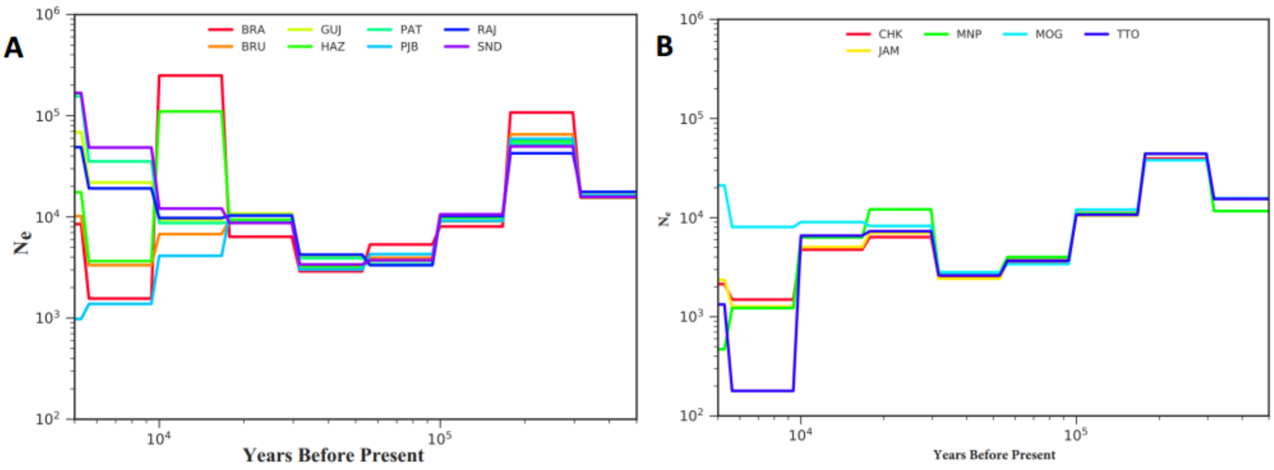


**Fig. S5.**

**Demographic histories of Pakistani populations and those speaking in Sino-Tibetan family languages. A.** Demographic histories for the Pakistani populations included in this study. Population size change histories are plotted from 500k ybp to 5k ybp. The populations are Brahui (BRA), Gujjar (GUJ), Pathan (PAT), Sindhi (SND), Burusho (BRU), Hazara (HAZ), Punjabi (PJB) and Rajput (RAJ). Please refer to **Table S1** for the details of these populations. **B.** Demographic histories for the SAS Sino-Tibetan speaking populations included in this study. Population size change histories are plotted from 500k ybp to 5k ybp. The populations are Chakma (CHK), Manipuri (MNP), Mog (MOG), Toto (TTO) and Jamatia (JAM). Please refer to **Table S1** for the details of these populations.

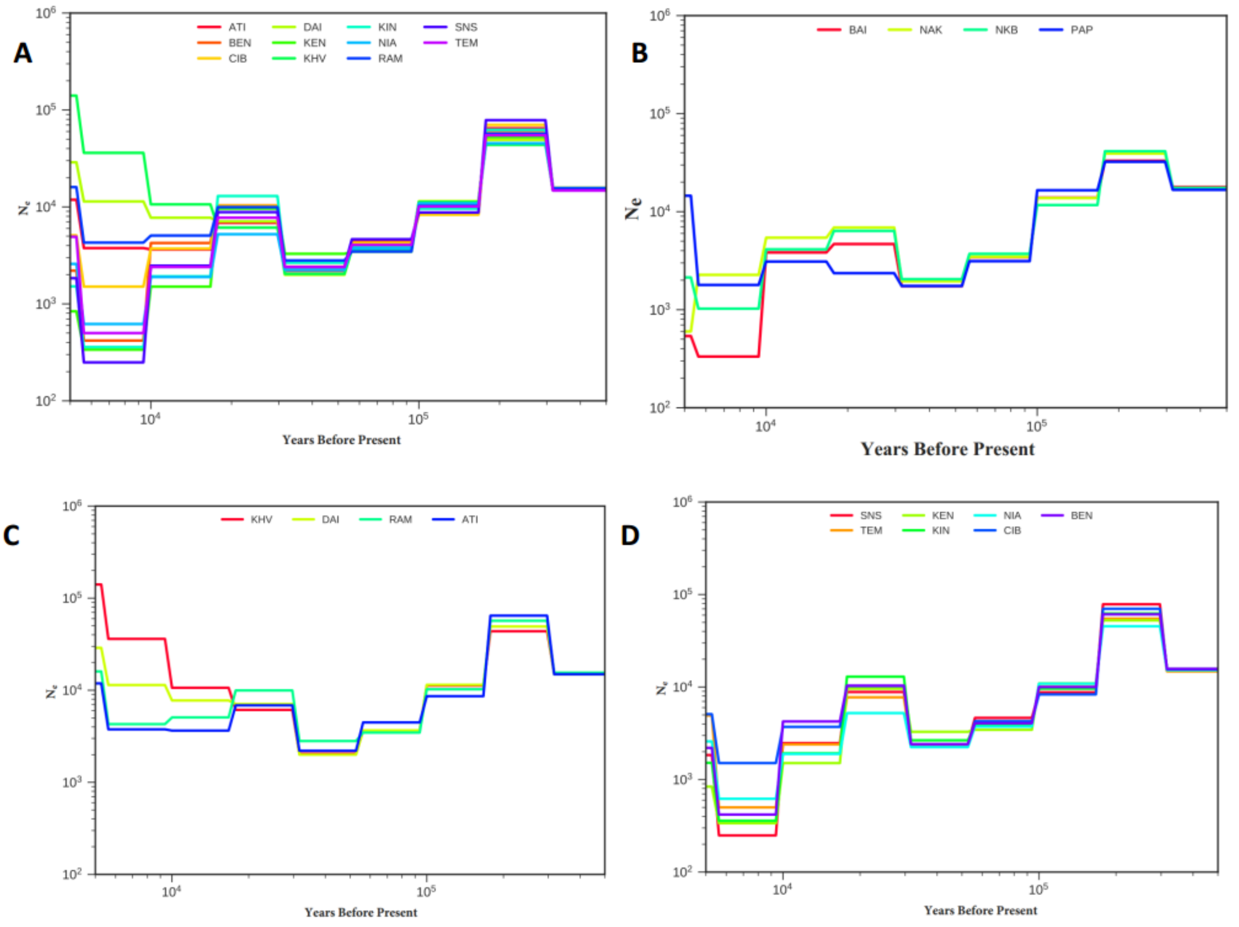


**Fig. S6.**

**Demographic histories for SEA (A) and OCE (B) populations included in this study.** Population size change histories are plotted from 500k ybp to 5k ybp. The populations are Ati (ATI), Dai (DAI), Kintak (KIN), Senoi Semai (SNS), Flores Bena (BEN), Kensiu (KEN), Austronesian (NIA), Temuan (TEM), Flores Cibal (CIB), Kinh from Vietnam (KHV) and Rampasasa (RAM). The SEA populations that went through a bottleneck ~10k ybp are shown in Figure **C** and the populations that experienced no change or an increase in Ne are shown in Figure **D**. The OCE populations in panel **B** are Nakani (NAK), Nakani Bileki (NKB), Baining (BAI) and Papuan (PAP).Please refer to **Table S1** for the details of these populations.

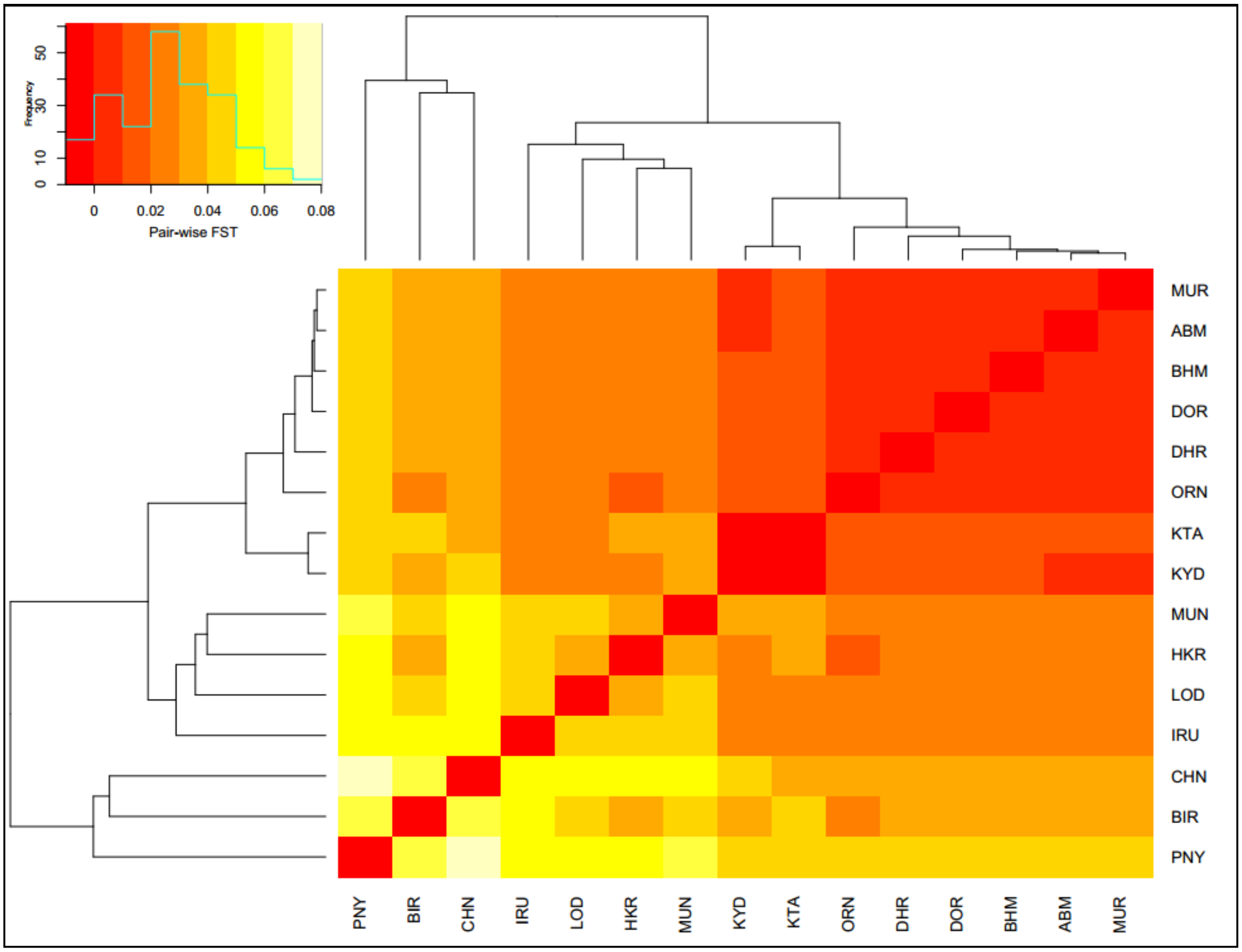


**Fig. S7.**

**$F_{ST}$ estimates for the Indian DR and AA speaking tribes.** The populations BIR, LOD, HKR and MUN are AA speakers and the rest are DR speakers.

| Population Code | Sample Size (N) | Population Name | Country | Region | Language Family | Lifestyle |
|---|---|---|---|---|---|---|
| **YRI** | 31 | Yoruba | Nigeria | Africa | Niger-Congo | Non Hunter Gatherer |
| **GBR** | 28 | Great Britain | UK | Europe | Indo European | Non Hunter Gatherer |
| **HAN** | 20 | Han Chinese | China | Northeast Asia | Sino Tibetan | Non Hunter Gatherer |
| **NAK*** | 7 | Nakanai | Papua New Guinea | Oceania | Austronesian | Hunter Gatherer |
| **NKB*** | 11 | Nakanai Bileki | Papua New Guinea | Oceania | Austronesian | Hunter Gatherer |
| **BAI*** | 19 | Baining | Papua New Guinea | Oceania | Baining | Hunter Gatherer |
| **PAP*** | 20 | Papuan | Papua New Guinea | Oceania | Papuan | Hunter Gatherer |
| **BIR*** | 20 | Birhor | India | South Asia | Austroasiatic | Hunter Gatherer |
| **HKR*** | 10 | Hill Korwa | India | South Asia | Austroasiatic | Hunter Gatherer |
| **LOD*** | 11 | Lodha | India | South Asia | Austroasiatic | Hunter Gatherer |
| **MUN*** | 2 | Munda | India | South Asia | Austroasiatic | Hunter Gatherer |
| **NIC*** | 6 | Nicobarese | India | South Asia | Austroasiatic | Hunter Gatherer |
| **IRU*** | 4 | Irula | India | South Asia | Dravidian | Hunter Gatherer |
| **BRA*** | 11 | Brahui | India | South Asia | Dravidian | Hunter Gatherer |
| **DHR*** | 10 | Dhurwa | India | South Asia | Dravidian | Hunter Gatherer |
| **DOR*** | 12 | Dorla | India | South Asia | Dravidian | Hunter Gatherer |
| **MUR*** | 10 | Muria | India | South Asia | Dravidian | Hunter Gatherer |
| **ABM*** | 11 | Abujmaria | India | South Asia | Dravidian | Hunter Gatherer |
| **BHM*** | 5 | Bison Horn Maria | India | South Asia | Dravidian | Hunter Gatherer |
| **ORN*** | 15 | Oraon | India | South Asia | Dravidian | Hunter Gatherer |
| **CHN*** | 9 | Chanchu | India | South Asia | Dravidian | Hunter Gatherer |
| **KYD*** | 11 | Kaya Dora | India | South Asia | Dravidian | Hunter Gatherer |
| **KNR*** | 17 | Konda Reddy | India | South Asia | Dravidian | Hunter Gatherer |
| **PNY*** | 11 | Paniya | India | South Asia | Dravidian | Hunter Gatherer |
| **KTA*** | 8 | Kota | India | South Asia | Dravidian | Hunter Gatherer |
| **IYE*** | 13 | Iyer | India | South Asia | Dravidian | Non Hunter Gatherer |
| **IYA*** | 6 | Iyengar | India | South Asia | Dravidian | Non Hunter Gatherer |
| **SOB*** | 9 | Saurashtra Brahmin | India | South Asia | Dravidian | Non Hunter Gatherer |
| **WBB*** | 10 | West Bengal Brahmin | India | South Asia | Indo European | Non Hunter Gatherer |
| **SRB*** | 14 | Saryupari Brahmin | India | South Asia | Indo European | Non Hunter Gatherer |
| **AGH*** | 17 | Agharia | India | South Asia | Indo European | Non Hunter Gatherer |
| **MHR*** | 19 | Mahar | India | South Asia | Indo European | Non Hunter Gatherer |
| **CHM*** | 6 | Chamar | India | South Asia | Indo European | Non Hunter Gatherer |
| **BAG*** | 3 | Bagdi | India | South Asia | Indo European | Non Hunter Gatherer |

| Population Code | Sample Size (N) | Population Name | Country | Region | Language Family | Lifestyle |
|---|---|---|---|---|---|---|
| **GAU*** | 9 | Gaud | India | South Asia | Indo European | Non Hunter Gatherer |
| **NAB*** | 4 | Nababuddha | India | South Asia | Indo European | Non Hunter Gatherer |
| **PJB*** | 5 | Punjabi | India | South Asia | Indo European | Non Hunter Gatherer |
| **RAJ*** | 14 | Rajput | India | South Asia | Indo European | Non Hunter Gatherer |
| **JAR*** | 12 | Jarwa | India | South Asia | Ongan | Hunter Gatherer |
| **ONG*** | 15 | Onge | India | South Asia | Ongan | Hunter Gatherer |
| **CHK*** | 11 | Chakma | India | South Asia | Sino Tibetan | Non Hunter Gatherer |
| **TTO*** | 11 | Toto | India | South Asia | Sino Tibetan | Non Hunter Gatherer |
| **MOG*** | 20 | Mog | India | South Asia | Sino Tibetan | Non Hunter Gatherer |
| **JAM*** | 9 | Jamatia | India | South Asia | Sino Tibetan | Non Hunter Gatherer |
| **MNP*** | 5 | Manipuri | India | South Asia | Sino Tibetan | Non Hunter Gatherer |
| **PAT*** | 17 | Pathan | Pakistan | South Asia | Indo European | Non Hunter Gatherer |
| **BRU*** | 10 | Burusho | Pakistan | South Asia | Indo European | Non Hunter Gatherer |
| **GUJ*** | 20 | Gujjar | Pakistan | South Asia | Indo European | Non Hunter Gatherer |
| **SND*** | 12 | Sindhi | Pakistan | South Asia | Indo European | Non Hunter Gatherer |
| **KHA*** | 11 | Khatri | Pakistan | South Asia | Indo European | Non Hunter Gatherer |
| **HAZ*** | 17 | Hazara | Pakistan | South Asia | Indo European | Non Hunter Gatherer |
| **DAI*** | 25 | Dai | China | Southeast Asia | Kra-Dai | Non Hunter Gatherer |
| **NIA*** | 15 | Austronesian | Indonesia | Southeast Asia | Austronesian | Hunter Gatherer |
| **CIB*** | 12 | Flores Cibal | Indonesia | Southeast Asia | Austronesian | Hunter Gatherer |
| **BEN*** | 11 | Flores Bena | Indonesia | Southeast Asia | Austronesian | Hunter Gatherer |
| **RAM*** | 20 | Flores Rampasasa | Indonesia | Southeast Asia | Austronesian | Hunter Gatherer |
| **SNS*** | 10 | Senoi Semai | Malayasia | Southeast Asia | Austroasiatic | Hunter Gatherer |
| **KIN*** | 19 | Kintak | Malayasia | Southeast Asia | Austroasiatic | Hunter Gatherer |
| **KEN*** | 9 | Kensiu | Malayasia | Southeast Asia | Austroasiatic | Hunter Gatherer |
| **TEM*** | 15 | Temuan | Malayasia | Southeast Asia | Austronesian | Hunter Gatherer |
| **ATI*** | 21 | Ati | Philippines | Southeast Asia | Austronesian | Hunter Gatherer |
| **KHV*** | 28 | Kinh | Vietnam | Southeast Asia | Austroasiatic | Non Hunter Gatherer |

**Table S1.**

**Description of the populations used in this study.** The populations marked with an '*' were included in the generalized liner model shown in Figure 2 and Supplementary Data S1.

| Time (ybp) | Median | Lower CI | Upper CI |
| --- | --- | --- | --- |
| 5623 | **2693.86** | **1966.89** | **3297.10** |
| 10000 | **3667.41** | **2656.57** | **4470.94** |
| 17783 | **792.78** | **-712.98** | **1745.65** |

**Table S2.**

**Summary of the bootstrap results.** The medians and 95% confidence intervals are shown for the three time points used for the comparison in **Fig. 1**.

| Time (ybp) | Median | Lower CI | Upper CI |
| --- | --- | --- | --- |
| 5623 | **2472.72** | **1339.11** | **3199.31** |
| 10000 | **2334.50** | **962.20** | **3971.485** |
| 17783 | **899.92** | **-754.28** | **1896.25** |

**Table S3.**

**Summary of the bootstrap results for SAS populations.** The medians and 95% confidence intervals are shown for the three time points used for the comparison in **Fig. S1**.
